# Metagenomic analysis of the honey bee queen microbiome reveals low bacterial diversity and Caudoviricetes phages

**DOI:** 10.1101/2023.08.29.555332

**Authors:** Lilian Caesar, Danny W. Rice, Alison McAfee, Robyn Underwood, David R. Tarpy, Leonard J. Foster, Irene L. G. Newton

## Abstract

In eusocial insects, the health of the queens – the colony founders and sole reproductive females – is a primary determinant for colony success. Queen failure in the honey bee *Apis mellifera*, for example, is a major concern of beekeepers that annually suffer with colony losses, necessitating a greater knowledge of queen health. Several studies on the microbiome of honey bees have characterized its diversity and shown its importance for the health of worker bees, the female non-reproductive caste. However, the microbiome of workers differs from that of queens, which in comparison is still poorly studied. Thus, direct investigations of the queen microbiome are required to understand colony-level microbiome assembly, functional roles, and evolution. Here we used metagenomics to comprehensively characterize the honey bee queen microbiome. Comparing samples from different geographic locations and breeder sources, we show that the microbiome of queens is mostly shaped by the environment experienced since early life, and is predicted to play roles in breakdown of the diet, and protection from pathogens and xenobiotics. We also reveal the microbiome of queens comprises only four core bacterial species, *Apilactobacillus kunkeei*, *Lactobacillus apis, Bombella apis* and *Commensalibacter* sp. Interestingly, in addition to bacteria, we show that bacteriophages infect the queen microbiome, for which Lactobacillaceae are predicted to be the main reservoirs. Together, our results provide the basis to understand the honey bee colony microbiome assemblage, can guide improvements in queen rearing processes, and highlight the importance of bacteriophages for queen microbiome health and microbiome homeostasis in eusocial insects.

**Importance:** The queen caste plays a central role for colony success in eusocial insects, as queens lay eggs, and regulate colony behavior and development. Queen failure can cause colonies to collapse, which is one of the major concerns of beekeepers. Thus, understanding of the biology behind the queen’s health is a pressing issue. Previous studies have shown the bee microbiome plays an important role in worker bee health, but little is known about the queen microbiome and its function *in vivo*. Here, we characterized the queen microbiome identifying for the first time present species and their putative functions. We show that the queen microbiome have predicted nutritional and protective roles in queen association, and comprises only four consistently present bacterial species. Additionally, we bring to attention the spread of phages in the queen microbiome, which increased in abundance in failing queens and may impact the fate of the colony.

## Introduction

Queen health in eusocial insects is a primary determinant for the success of the colony. Eusocial insects live in groups with cooperative care of juveniles, overlap of generations, and reproductive division of labor. Exemplars of insect societies are ants, wasps, and bees from the order Hymenoptera, the largest and most well-known animal group with eusocial species. In the eusocial insects, the reproductive female caste (queens) is responsible for laying eggs and regulating the colony’s behavior and development. After mating, queens spend almost their entire life, which can reach several decades, inside the colony being fed by specialized workers. They are among the most fecund terrestrial animals; queens from some insect species can lay ∼20,000 eggs per day (1), which may explain the success of some eusocial insect genera. The importance of queens also extends beyond their reproductive role. Queens maintain the colony homeostasis by managing the behavior of other colony members, such as attracting workers or inducing submissive behavior, modulating aggression, or inhibiting production of new queens through pheromone signals (2, 3, 4, 5, 6).

The importance of queens for colony success becomes even more evident with the example of honey bees (*Apis mellifera*). In addition to the significance of honey bees for general ecosystem services in their natural range, managed colonies contribute billions to the agricultural economy annually in the US, due to their pollination services (7). However, annual losses of honey bee managed colonies have reduced reliability on them and affected food security. Queen failure and premature supersedure are consistently reported as the leading contributing factors for colony mortality (8). Honey bee queens can live three to four years, but recently, their diminished longevity requires the replacement of the queens almost every year, a practice also used preventively (9). The causes behind failing queens are still poorly understood, but it may include problems with development, insemination success, infection by parasites, exposure to xenobiotics, and adverse temperatures (10, 11). Interestingly, the microbiome of insects, including honey bee foragers, is also known to modulate the host response to some of these stressors. For example, the worker gut microbiome can protect the host from parasites (12), and parasites can also shape the gut microbiome due to the infection (13), ultimately impacting colony fitness. Thus, the microbiome is an important trait that should be included in studies on the health of eusocial insect queens, and *Apis mellifera* is an excellent model system for these investigations.

Most of what is known about the microbiome of honey bees and its role in colony health comes from investigations on worker bees, primarily foragers – the non-reproductive females that frequently leave the colony to collect pollen and nectar. Honey bee workers have a taxonomically simple and consistent gut microbiome among colonies around the world, comprised of the core members *Bifidobacterium*, *Snodgrassella*, *Gilliamella*, and two groups of *Lactobacillus,* Firm-4 and Firm-5. This microbial community impacts host health in multiple ways (see review, 14), and its acquisition and transmission occur mostly through interactions with individuals from the colony and the hive environment (15). Dysbiosis in this system is strongly correlated with poorer worker fitness and is characterized by shifts in the load of core microbes and the spread of opportunistic bacteria (16). More recently, fungi and bacteriophages community characterization have also been included in studies of the bee microbiome (17, 18). The role that phages play in molding the microbiome and impacting host health has been shown in unrelated model systems (19, 20, 21, 22), but in honey bees their diversity and potential effects are under studied.

Curiously, there is evidence that honey bee queen microbes are distinct from those of workers from their own colony (23). Thus far, based on 16S rRNA amplicon-based studies, the honey bee queen microbiome appears to comprise mostly bacteria from the genera *Lactobacillus*, *Bombella* and *Commensalibacter*. However, the queen gut microbiome is not as consistent as observed for worker bees, and the relative abundance of the associated bacteria is quite variable among queens from different colonies (23, 24, 25). In addition to the natural microbiome variability among queens, controlled experiments have shown that both age and early-bacteria colonizers coming from social interactions or rearing protocols also lead to differences in their microbiome (24). A metagenomic approach, enabling species-level characterization and access to genomic information, could improve our understanding of the queen microbiome assemblage and its role in queen health.

Here we sequenced the metagenomes of 18 queen gut samples from the US and Canada to deeply characterize their microbiome. These queen samples have associated metadata from previous studies (26, 27), including both failing and healthy queens. We describe the queen gut microbial community, from bacteria to phages, and investigate the most important factors shaping it. In addition, we characterize the microbiome at a functional level, and recover metagenome-assembled genomes to identify, at the species level, the candidate core microbiome.

## Results

### The honey bee queen microbiome is dominated by bacteria of the Acetobacteraceae and Lactobacillaceae families

We sequenced a total of 18 queen gut metagenomes, ranging from 7 to 33 million paired-reads per sample after quality trimming (Table S1). First, we asked which microbes were present in the community associated with the honey bee queen gut. We mapped trimmed reads against 18S rRNA and 16S rRNA databases for taxonomic identification of fungi and bacteria, respectively. As expected, based on the paucity of fungal reads recovered from other studies, no reads mapped to 18S rRNA; on the other hand, reads mapped to bacteria from at least five families, mostly Acetobacteraceae and Lactobacillaceae (Fig. 1A). At the genus level, the queen microbiomes varied extensively in the abundance of Acetobacteraceae (*Bombella* or *Commensalibacter*), and Lactobacillaceae (*Apilactobacillus* or *Lactobacillus*) (Fig. 1A). Other bacteria, known to often be part of the worker bee gut microbiome, were also present in some of the queen samples, such as *Bombilactobacillus*, *Bifidobacterium* and *Frischella*. The geographical location from which queens were sampled (permanova; *City*, bacterial family, p = 0.046, R2 = 0.304, F = 2.165; bacterial genus p = 0.045, R2 = 0.252, F = 1.667; *State*, bacterial family, p = 0.047, R2 = 0.115, F = 2.270; bacterial genus, p = 0.039, R2 = 0.121, F = 2.286) and the year of sampling (which is conflated with the geographic location, see Table S1), were two of the factors explaining the differences between queen microbiomes both at the family and genus level. Additionally, the queen source – that is, the breeder who provided the queen – was also a factor explaining microbiome differences across samples (permanova; bacterial family, p = 0.040, R2 = 0.421, F = 2.488, bacterial genus, p = 0.039, R2 = 0.357, F = 1.850). Queen source, although influenced by genetics of the stocks, could also be influenced by environmental differences since the queen rearing environment and protocol used by the beekeeper could impact queen microbiomes. Importantly, in our study, the location of these samples (*State* and *City*) is also confounded by queen source, as beekeepers from the same area generally had the same queen suppliers (Table S1).

**Figure 1:**
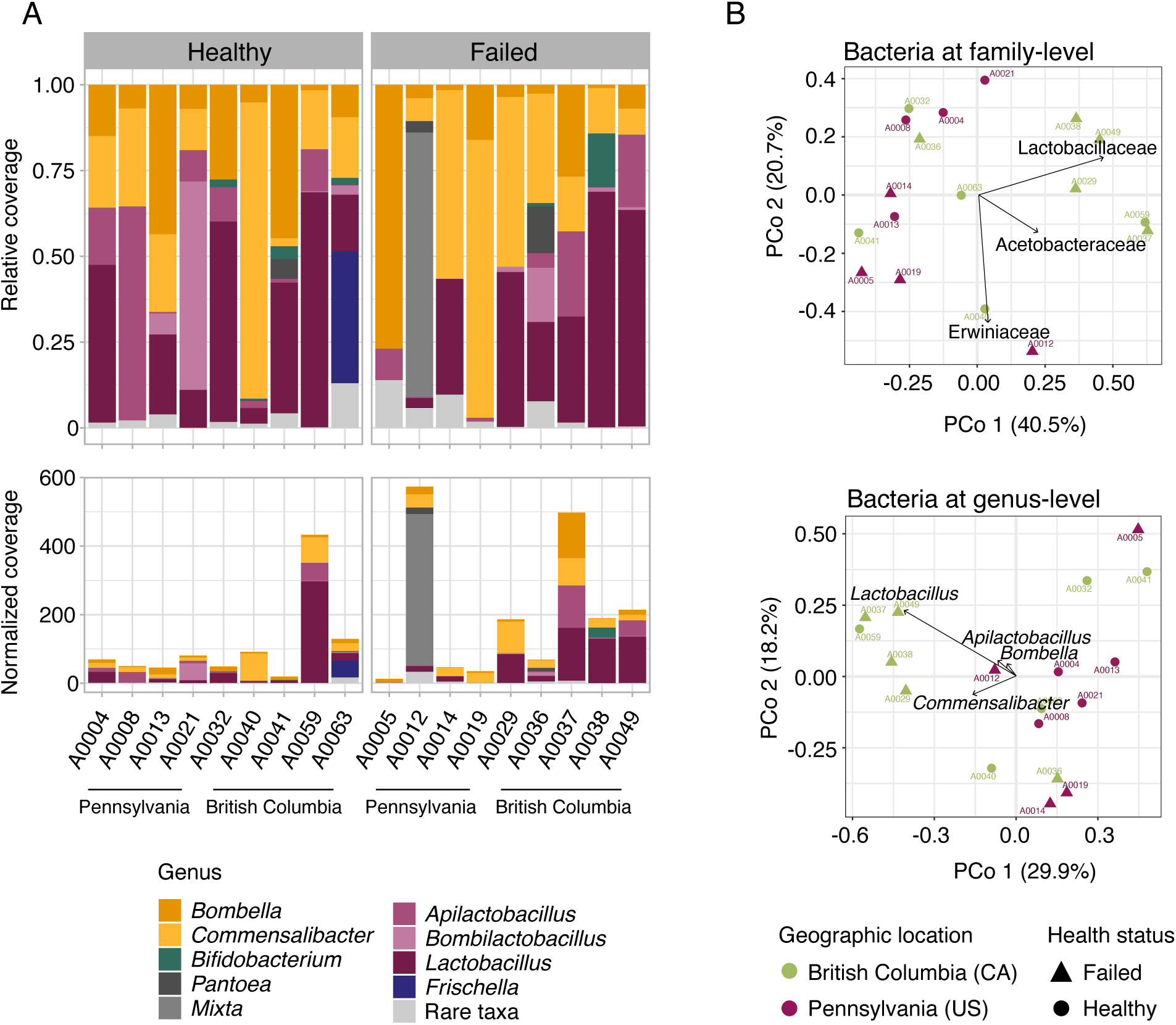
The microbiome of honey bee queens varies extensively in the abundance of the two most prevalent bacterial families, Acetobacteraceae and Lactobacillaceae (A) Relative (above) and absolute normalized (below) coverage of bacteria genera comprising the microbiome of healthy and failing honey bee queens from two different locations, Pennsylvania (US) and British Columbia (CA) Color shades represent bacteria families; Acetobacteraceae (yellow), Bifidobacteriaceae (green), Erwiniaceae (gray), Lactobacillaceae (purple), Orbaceae (blue) (B) Principal Coordinate Analysis (PCoA) showing the distribution of queen samples based on the dissimilarity of relative proportions of the microbial communities at family-(above) and genus-level (below) Arrows highlight the bacteria with the strongest contribution to the ordination, and each axis shows the percentage of variation explaining

Due to the effect of the environment on the queen microbiome composition, differences in the microbiome of healthy or failing queens were only observed depending on their geographical location (permanova; interaction between *State* and *Status*, p = 0.047, R2 = 0.121, F = 2.395). This interaction effect was observed at the family level; the queen core microbiome families, Lactobacillaceae and Acetobacteraceae, were among the taxa that explain most of these differences among the samples and their clustering in the Principal Coordinate Analysis, PCoA (Fig. 1B). But also, Erwiniaceae, a non-core bacteria family of the honey bee microbiome, was mostly found in failing queens and also had an important contribution to the PCoA topology (Fig. 1B). Interestingly, with the normalization of the bacteria coverage by the library size, we can also observe that failing queens, mostly from British Columbia (CA), have higher loads of bacteria per gut sample (Fig. 1A).

### Queen genetic background is not directly correlated with microbiome composition or health status

Because the queen source – that is, the breeder who provided the queen – was one of the factors explaining microbiome composition, we decided to further investigate if the queen’s genetic background could be a direct predictor of its gut community. Honey bee queen population structure was assessed from genotype likelihoods of 7,342,540 polymorphic sites. Both population structure plots showed a clear genetic differentiation among the honey bee queens from Pennsylvania (US) and British Columbia (CA); although the admixture plot suggests some gene flow between populations (Fig. 2A). We also tested higher numbers of ancestral populations and there is no clear population structure for queens from the same apiary or queen company supplier (Fig. S1). The principal-component analysis (PCA) also cleanly delineates the samples based on State factor and interestingly, the samples with mixed ancestry are those at the edges of each cluster, closer to the center of PC1 (Fig. 2B). Finally, we used a Mantel test to compare similarity/dissimilarity of queen genetic background and microbiome composition. The bacterial families or genera were not correlated with pairwise genetic distance among queens (bacterial family, R = 0.04113, p = 0.3906; bacterial genus R = 0.1676, p = 0.1152) or covariance matrix of queen genotype likelihoods (bacterial family, R = 0.05412, p = 0.2925; bacterial genus R = −0.04525, p = 0.727).

**Figure 2:**
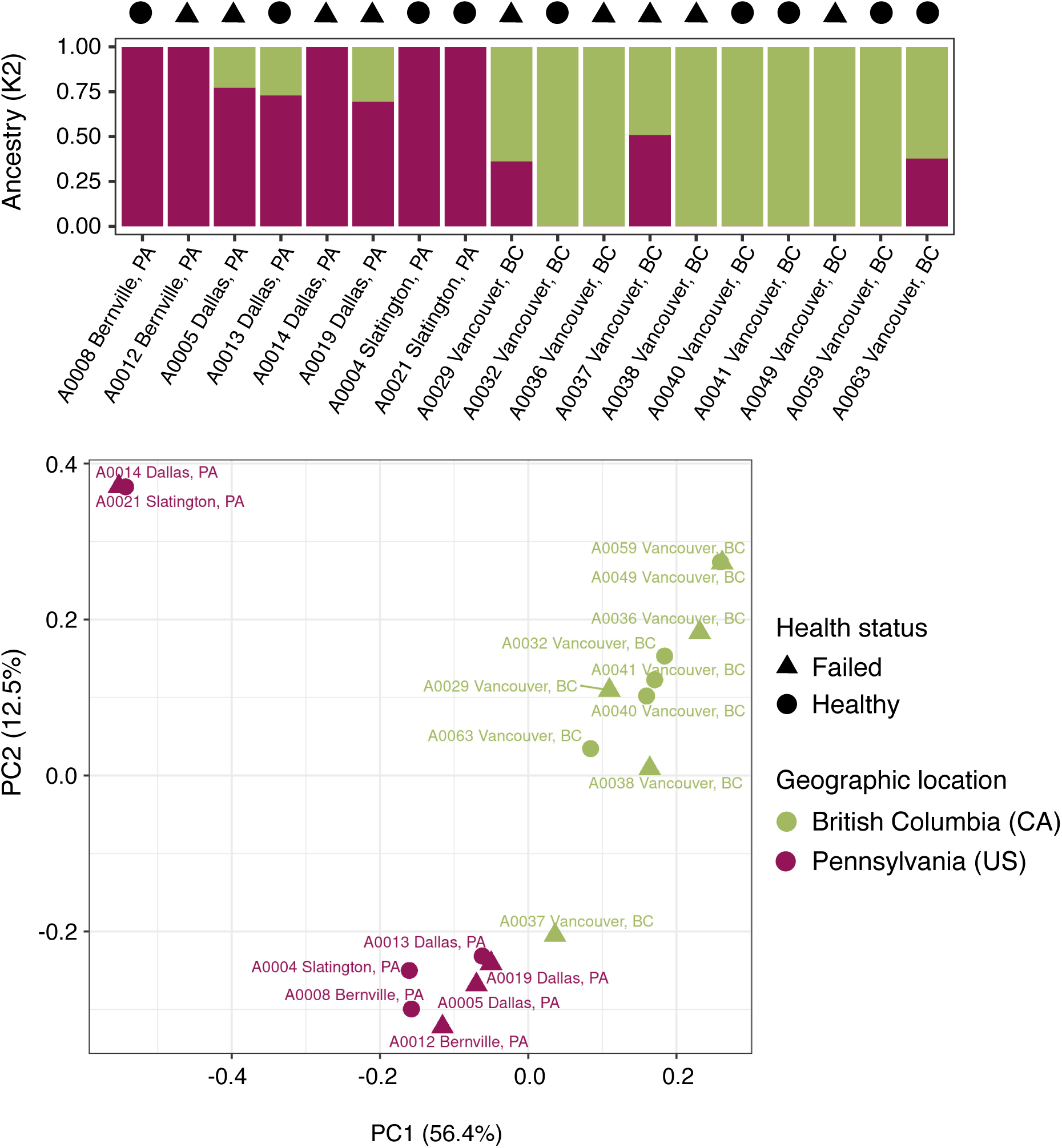
Genetic background of the honey bee queens groups them in a population for each general geographical location, and not by health status or microbiome composition Above, the bar plot resulting from the SNP admixture analysis for K2 ancestral populations (y-axis shows the proportion of each ancestral population), clustering queens from Pennsylvania (PA) and British Columbia (CA) Below, the Principal Component Analysis (PCA) of the genotype likelihood covariance matrix shows the same population structure, separated in PC 1 that explains 56 4% of the total variation

### The candidate core microbiome of queens comprises four bacterial species

We recovered metagenome-assembled genomes (MAGs) of the core honey bee queen microbiome members. The co-assembly of all trimmed reads generated 177,846 contigs > 500 nt (Table S2), which were then grouped into bins. In total, 8 bacterial MAGs with partial (<50%) to nearly complete (>90%) genomes were recovered and placed into a phylogenetic context to predict species (Table S3, Fig. 3, Fig. S2). Considering that MAGs will be retrieved from the abundant species in metagenomes (present in more samples, with more reads), among them we should be able to find the core members of the queen microbiome. Core microbiome members should be present in all healthy queens and some of the failing queens, while non-core members would be present only in a few queen samples. Single-copy core genes were used to estimate the coverage of MAGs, revealing that *Bombella apis*, *Commensalibacter* sp., *Apilactobacillus kunkeei* and *Lactobacillus apis* were present in all queen samples (with the exception of *L. apis* which was absent in one failing queen; Fig. 3, Table S4). In 11 out of 18 samples, these four bacteria species represent more than 50% of the microbiome (Table S4). Only the sample A0012, a failing queen, had an important reduction of the core microbiome (9%) and an increase in the abundance of an Enterobacteriaceae. This opportunistic bacterium is part of the non-core MAGs, which includes the Lactobacillaceae *L. panisapium*, the Bifidobacteriaceae *Bifidobacterium apousia*, the Orbaceae *Frischella perrara* and the Enterobacteriaceae *Tenebrionicola*-like (Fig. S2).

**Figure 3:**
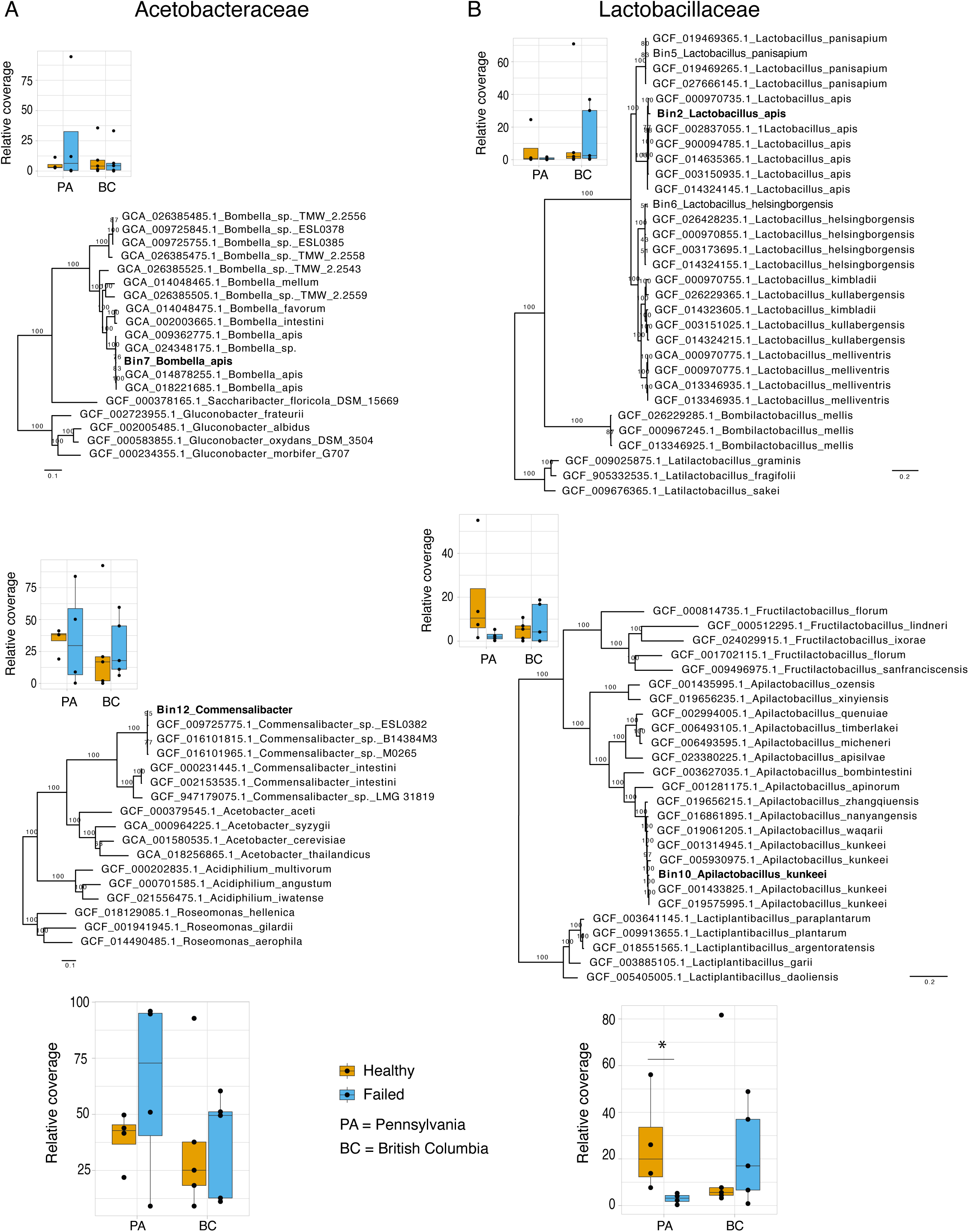
The core microbiome of honey bee queens comprises only four species. (A) Phylogenies inferred with maximum likelihood for the metage-nome-assembled genomes (MAGs) of the two core Acetobacteraceae. Above, Bombella apis and relatives, inferred from the alignment of 927 single-copy orthologs (SCO). Below, Commensalibacter and relatives, inferred from the alignment of 708 SCO. (B) Phylogenies for the MAGs of the two core Lactobacillaceae. Above, Lactobacillus apis and relatives, inferred from the alignment of 80 SCO. Below, Apilactobacillus kunkeei and relatives, inferred from the alignment of 292 SCO. (A, B) On the side of each phylogeny is the relative coverage of the MAG in queens with different health status and geographical location. Below the phylogenies of each bacterial family is the same plot configuration, but with the coverage sum of both MAGs. Significant statistical difference was only observed when comparing the abundance of core Lactobacillaceae among queens with different health status in Pennsylvania; there is a reduction of their abundance in failing queens (Wilcoxon rank test, p = 0.028, *).

We also used these MAGs to then ask if a shift in microbial community, i.e., dysbiosis, was observed in the abundance of the core species from the microbiome of failing queens. Although we can observe a tendency for a shift in species proportions when comparing queens with different health statuses, variance of relative coverage within the group was high, and there was no significant difference and no shift pattern in their proportions in failing queens across both locations (Fig. 3). At the family level, however, *Lactobacillus* in the state of Pennsylvania had a significant decrease in their proportion in failing queens (Wilcoxon rank test, p = 0.028), while for Acetobacteraceae this difference is not statistically significant (*t-*test, p = 0.319, df = 6).

### Functional properties of the queen gut microbiome

To characterize the potential role of the microbiome in the queen host, and interactions with and within the microbial community, we predicted protein functions from all bacterial contigs – thus avoiding biases due to MAG completion status or non-binned partial bacteria genomes which may also have proteins with functional relevance. From all bacterial contigs, 23,833 proteins were predicted and 12,910 (54%) were assigned to KEGG ortholog categories. Presence and absence of proteins in each sample was used to compare them in a clustered heatmap, where all functional categories in the queen microbiome are shown (Fig. 4). Overall, there was no broad functional category missing or overrepresented in samples, so no clustering by microbiome functional profile was observed for samples of the same geographic location (Pennsylvania vs. British Columbia) or health status. Cluster 2 (C2, Fig. 4), however, includes the samples with more abundant non-core bacteria. In addition to proteins with general cell function, the main categories present were metabolism of cofactors and vitamins, energy metabolism, carbohydrate metabolism, and amino acid metabolism. Protein counts from the energy metabolism category confirm that the microbes associated with honey bee queens mostly use oxidative phosphorylation for energy production (Fig. S3E). These microbes also harbor genes involved in the degradation of xenobiotics (e.g., cytochrome P450), the biosynthesis of antibiotics (e.g., monobactam and streptomycin), and biofilm formation pathways which may play an important role in community regulation (Fig. S3M, B and D, respectively). While genes for lysine biosynthesis were annotated in all samples, only sample A0012, a failing queen with depleted core-microbiome and increased abundance of the Enterobacteriaceae *Tenebrionicola*-like, have an increase in proteins involved in degradation of this amino acid, known to be able to buffer poor bee nutrition (Fig. S3A; 28).

**Figure 4:**
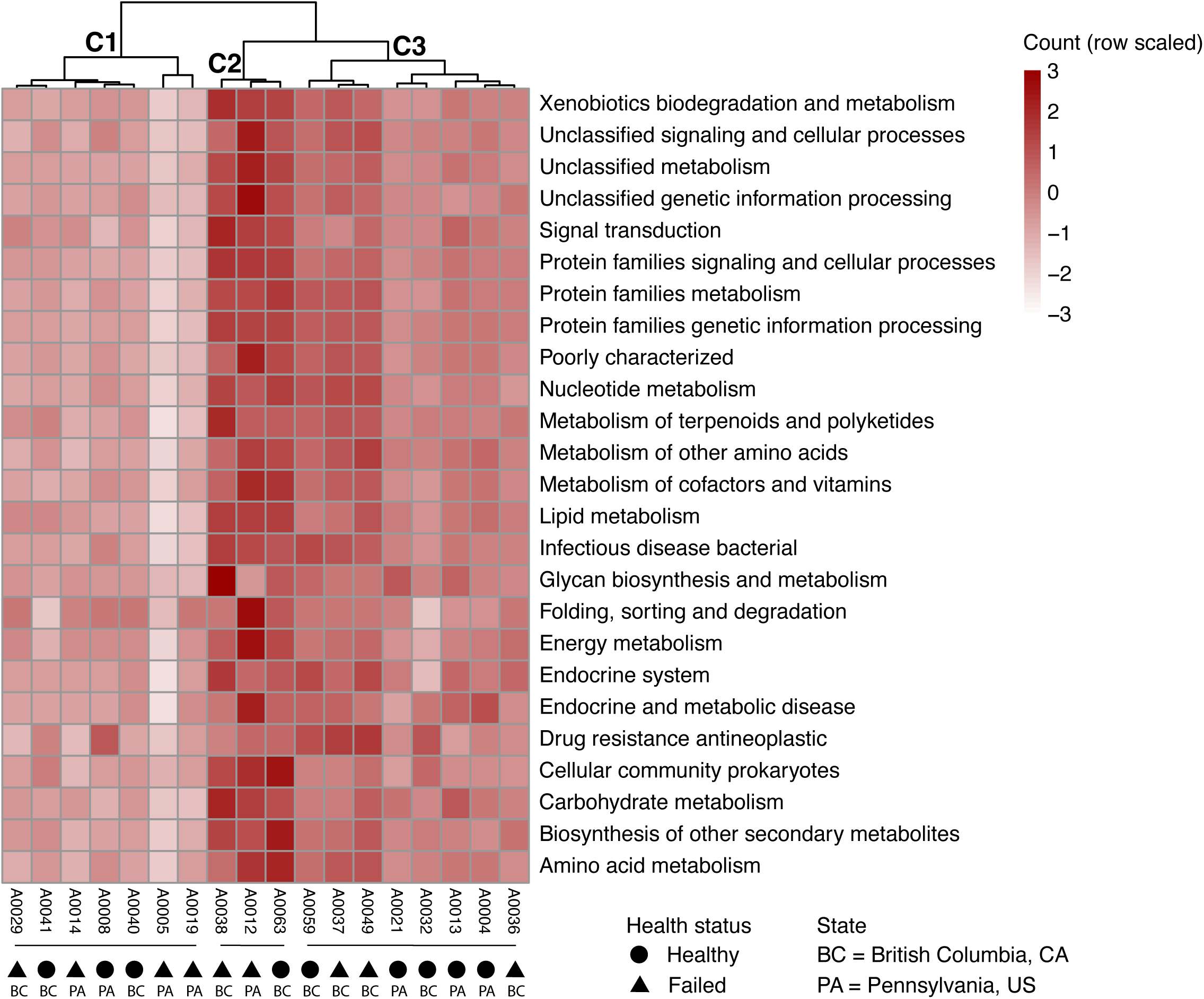
The microbiome functional profile suggests an important role for honey bee queen health and homeostasis. The heatmap shows scaled protein counts per KEGG category (rows), which include proteins involved in xenobiotics biodegradation and metabolism, amino acid metabolism biosynthesis (e.g., lysine) or biosynthesis of secondary metabolites (e.g., antibiotics). Samples (columns) were hierarchically clustered. Three main functional groups were formed (C1, C2, C3), but they do not correlate with any of the variables tested in our investigations, such as health status or geographic location. Cluster 2, however, has an increase (dark red) in the representation of all KEGG categories due to the relative abundance of non-core bacteria in their microbiomes.

### Bacteriophages of the class Caudoviricetes compose the queen microbiome and are found in higher abundance in failing queens

In addition to the search for fungi and bacteria in the microbiome of queens, we also wondered if bacteriophages, important modulators of microbial communities, could also compose the microbiome of the honey bee queens. A total of 74 phage vOTUs were retrieved, including 3 lysogenic and 71 putative lytic phages (Table S5). All the phages with predicted genes were classified taxonomically as Caudoviricetes based on the similarity with viral marker genes in the database. Among the lysogenic phages, one is integrated in the genome of *Lactobacillus panisapium*, and the other two are integrated in the genome of the Enterobacteriaceae, *Tenebrionicola*-like. To predict putative hosts for the lytic phages we first conducted nucleotide similarity searches against CRISPR-cas arrays predicted in our MAGs.Genomes of *B. apis, L. apis, L. panisapium* and *Tenebrionicola*-like have CRISPR-cas arrays with one to seven spacers per array (List S1), but none of the lytic phages from the metagenomes matched these spacers. However, we still were able to predict putative hosts of 21 lytic phages based on the other approaches used, including matches with other CRISPR spacers databases, BLASTs and k-mer compositional analyses (Fig. 5, Table S5). The lytic phages were predicted to infect mostly Lactobacillaceae and Enterobacteriaceae.

**Figure 5:**
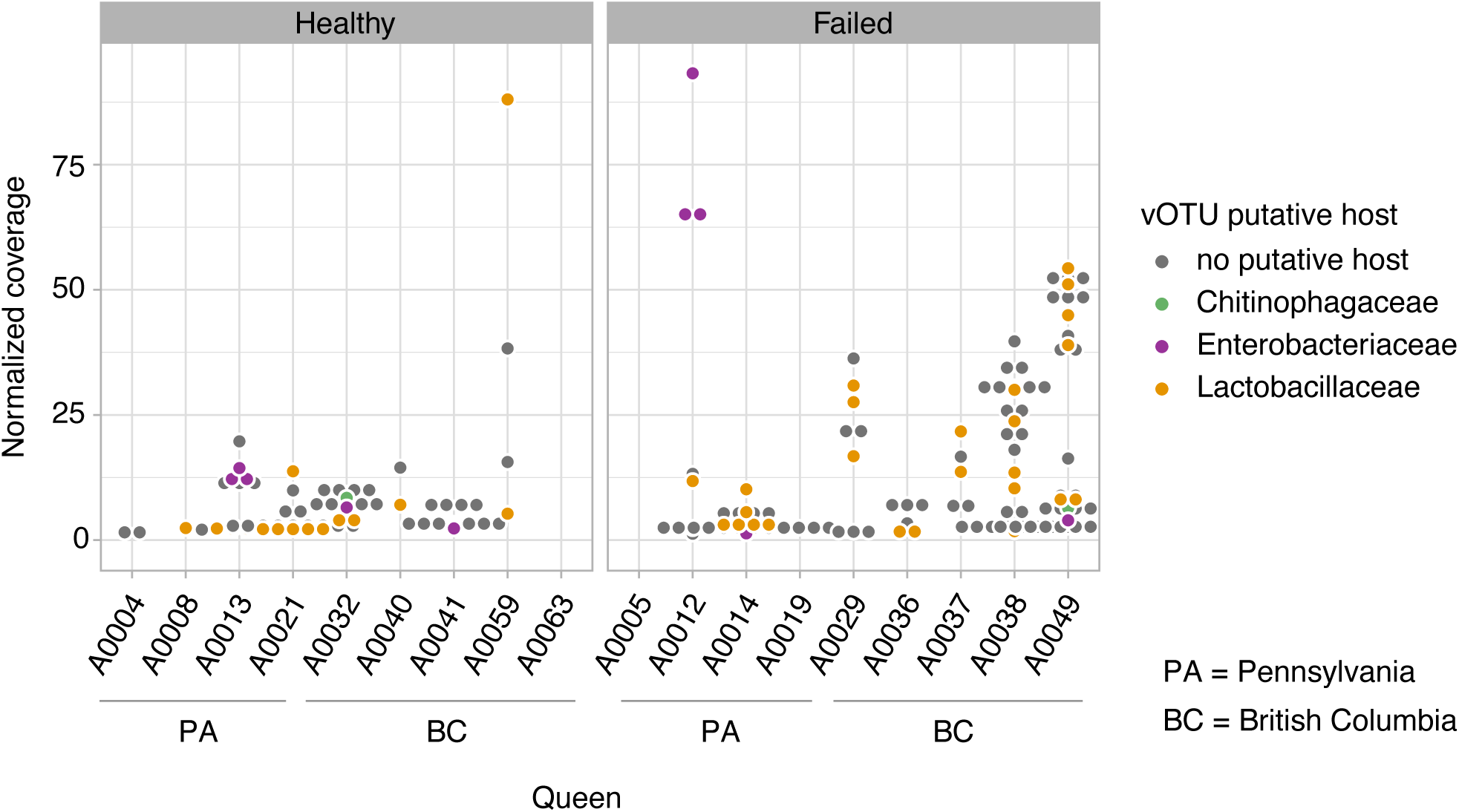
Caudoviricetes phages spread opportunistically in the microbiome of failing queens. The coverage of phage vOTUs (dots, with a coverage >1) are shown for each queen sample, which was higher in failing queens (Wilcoxon rank test, p = 0.029; PA, p = 0.032; BC, p = 5.12e-5). The color of the dot indicates the host where its genome is integrated or its the putative host.

Differences in the abundance of phages among the samples were significant for all factors being tested, including geographic location (Kruskal-Wallis rank test; p = 0.005, df = 3), age (Wilcoxon rank test; p = 8.41e-02), or queen source (Kruskal-Wallis rank test; p = 0.005, df = 4). Interestingly, the queen’s health status was one of most significant factors in terms of differences in phage abundance (Wilcoxon rank test; p = 0.029; British Columbia, p = 5.12e-05, Pennsylvania, p = 0.032). Also, since phages need a host, we could expect a consistent increased abundance of phages in the samples with increased abundance of their hosts. Following this rationale, we used a correlation analysis to potentially find putative hosts for phages with no putative host identified with the previous methods. As expected, all significant correlations among phages and bacteria were positive correlations, and most bacteria co-occurred precisely with phages predicted to infect their family, such as Lactobacillaceae and Enterobacteriaceae. The result of this analysis indicates some phages that may be candidates to infect three members of the core microbiome, *B. apis* (e.g., k141_169038), *A. kunkeei* (e.g., k141_368347) and *Commensalibacter* (e.g., k141_456794; Fig S4), although these bacteria also had significant, but weaker, correlations with Lactobacillaceae-putative phages.

## Discussion

Investigations on the microbiome of eusocial insect queens are not as frequent as those on worker castes, despite the key role of the queen for superorganism fitness. However, studies conducted so far have revealed that the microbiome of queens – including species of termites, ants and bees – usually differs from that of their workers (23, 29, 30). The lack of direct correlation between worker and queen microbiomes reinforces the importance of queen microbiome characterization to understand colony-level microbiome assembly, functional roles and evolution. Honey bees, especially, are economically relevant and have been important models for microbiome investigations; amplicon-based characterizations of the queen microbiome have shown that it is mostly comprised of Lactobacillaceae and Acetobacteraceae (23, 24, 25). Here, using a metagenomic approach, we showed that the microbiome of honey bee queens (I) varies considerably in the abundance of these two bacterial families, with the environment having an important impact on these differences, (II) the microbiome comprises four core bacterial species, and is predicted to have a protective and nutritional role, and (III) lysogenic and lytic Caudoviricetes phages are part of the microbiome, for which core Lactobacillaceae may be important reservoirs and spreaders.

Previous studies on bee workers and queens have shown that, to some extent, their genetic background influences microbiome composition. In workers these results were obtained by comparing the microbiome of *A. mellifera* subspecies, isolated from each other and inbred since 1980 (31). The evidence for the effect of queen genetic background on microbiome composition was shown by comparing two species of *Apis* (*A. cerana* and *A mellifera*), which have their last common ancestor dated to approximately 6 mya (32, 33). In the current study, we did not observe the same trends comparing queen genotypes of the same *A. mellifera* subspecies, meaning the genetic differences among the queens were not enough to explain differences in the microbiome (Fig. 2). Instead, we found that the queen microbiome composition is shaped by the environment, i.e., geographic location and queen breeder source. For worker bees it is known that the environment landscape plays an important role on the microbiome composition (34). The effect of breeder source on queen microbiome composition was also observed in another study with honey bee queens (25) and, together with the lack of correlation between queen genetic background and microbiome composition, suggests a role of priority effects (arrival order and/or timing) on queen microbiome assembly. The effect of the environment on microbiome composition may also have masked differences in the microbiome of healthy and failing queens (Fig. 1). Dysbiosis can shift the microbiome to a different end community composition depending on previous colonizers and the factors causing the shift (16). Thus, to characterize dysbiosis patterns in failing queens an experimental design controlling for environmental effects would be important in future investigations. Regardless, with our approach we were able to observe a typical dysbiotic pattern in some of the failing queens, which was an increase in bacteria abundance and an increase in the proportion of non-core microbiome members. However, the changes observed in the microbiomes of the failing queens in our study could be a consequence of weak queens, and not the direct cause of queen failure.

Our findings reveal that the microbiome of queens is very constrained, composed of only four core bacterial species: *Bombella apis, Commensalibacter sp., Apilactobacillus kunkeei* and *Lactobacillus apis* (Fig. 3). This represents even fewer bacterial members than the known simple core microbiome of worker bees, comprising five phylotypes with multiple species (35, 36). Additionally, no fungi were detected in our queen samples, which is in contrast to previously published amplicon sequencing-based investigations with queens and worker bees (17, 37, 38, 39). Importantly, this difference could be due to the fact that amplicon sequencing will detect extremely low abundance templates resulting from the honey bee diet.

Queens are the longest-lived members in a colony, they are larger than workers, and they have well-developed ovaries, thus their physiology itself already represents a different niche for bacteria to colonize. Additionally, the highly specialized diet of queens since larval development to adult life, which is primarily composed of royal jelly, may have facilitated the selection of a small number of core members. Queens are genetically identical to their sisters that became workers, i.e., female larvae are bipotent and can equally develop into queens or workers depending on their larval diet (royal jelly vs. worker brood food). Royal jelly is composed of water, proteins, sugars, and lipids, but it is also very viscous, acidic, and imbibed with antimicrobial peptides, such as royalisin, jelleines, apismin, royalactin and fatty acids, that together confer the antimicrobial role of the royal jelly (40, 41). For example, royal jelly can inhibit the growth of bee pathogenic bacteria, such as *Melissococcus plutonius* and *Paenibacillus alvei* (42). Interestingly, royal jelly does not inhibit all bacteria; a recent study showed that *Bombella apis*, one of the core members of the queen microbiome, can withstand and even replicate in royal jelly (28). In this same study, however, *Apilactobacillus kunkeei*, did not show the same ability. In other host model systems where the microbiome is also composed of Acetobacteraceae and Lactobacillaceae, such as in *Drosophila*, the colonization of these bacterial families is enabled due to cross-feeding; *Acetobacter pomorum* uses the lactate produced by *Lactobacillus plantarum* to supply amino acids that are essential to *L. plantarum* (43). All the members of the queen microbiome can be isolated and cultivated in the laboratory, and future experiments could test if cross-feeding is behind the presence of these core bacteria in such an antimicrobial environment. It is also important to note that the royal jelly is produced, secreted, and offered to the queen by nurse bees, meaning the selection for these royal-jelly resistant microbes is likely already occurring in the nurse bee hypopharyngeal glands. Moreover, it is conceivable that queens have different needs with regard to nutrition compared to workers, and thus they may require different nutritional symbionts. In fact, our functional analysis has shown that the microbiome of queens is equipped with genes involved in protein and carbohydrate metabolism (e.g., peptidases, lipid biosynthesis, fructose and mannose metabolism; see Fig. 4 and Fig. S3), likely supplementing primary components of the host’s diet.

With our metagenomic approach, we were able to show for the first time that Caudoviricetes, a class of tailed bacteriophages, are part of the queen gut microbial community (Table S5). Previous studies have shown that Caudoviricetes are the main phages in the microbiome of honey bee workers (18, 44, 45), but the lack of similarity in the gut bacterial community of these two female castes had left open the question about the phages present in the queen microbiome. In the worker gut the main phage hosts are the core bacteria *Bifidobacterium*, *Gilliamella* and *Lactobacillus,* and the non-core *Bartonella* (18, 44, 45). Among these hosts, *Lactobacillus* is the only genus that is also found in the queen microbiome and, interestingly, we show here it is the queen gut bacterium hosting the majority of phages (Fig. 5). Importantly, however, for many phages, host prediction is difficult to impossible (18, 44, 45); here 77% of the phage sequences did not have a host predicted. However, we indeed predicted CRISPR-Cas systems with spacers in the *Bombella apis* MAG, suggesting a history of infections by phages (List S1). Also, phage sequence distribution was consistent with the change in the microbiome due to environmental effects, and phages were also more abundant in the failing queens (Fig. 5). This result points out that dysbiosis, usually detected via shifts in bacterial composition, can also be characterized by phage production and spread in the microbiome. We observed that one of the most abundant phage sequences in the queen microbiome is associated with a non-core bacterium, an Enterobacteriaceae, putting the native queen microbiome at risk. Once phages are spread in the microbiome, they can not only directly kill their hosts, but may also trigger immunological responses or even foster gene transfer between cells (46). The inherent complexity of a eusocial insect colony poses a challenge in investigations on the health of queens and, ultimately, of the colony (47). For future studies, a longitudinal approach could improve the resolution of the ecological interaction feedback between phages and bacteria over time, and its impact on queen health or failure.

## Methods

### Honey bee queen samples

Honey bee queens were sampled in 2018 from colonies located in Pennsylvania, USA, and in 2019 from colonies in British Columbia, Canada. The samples from both locations are part of previously published datasets (26, 27), in which the queens were classified according to their health status. Queens were scored as healthy when they showed no sign of supersedure cells, no drone brood in worker cells, no signs of disease, and had strong worker populations. Failing queens exhibited one or more of the following: drone brood in worker cells, spotty brood pattern, weak colony population, and supersedure cells. In addition to the health status, these samples have associated metadata that was used for the analysis, including queen age, source, ovary mass and size, and management strategy (Table S1). All samples were stored at −70°C until dissection and further use.

### Sample processing and shotgun sequencing

All queens were dissected on ice with 70% ethanol sterilized tools by gently pulling on the last abdominal tergite. DNA extraction was performed by a microbiome analytics company, Microbiome Insights™, using the Qiagen MagAttract PowerSoil DNA KF Kit optimized for the Thermo Scientific KingFisher robot. Quality and quantity of DNA was checked on an Agilent 2200 TapeStation, and a total of 18 samples with a DNA Integrity Number (DIN) >6 were submitted for library construction and metagenome sequencing at the biotechnology company SeqCenter, Pittsburgh, PA. Libraries were prepared using the Illumina DNA Prep kit and IDT 10bp UDI indices, and sequenced on an Illumina NextSeq 2000, producing 2x151bp reads. Demultiplexing, quality control and adapter trimming was performed with bcl-convert (v3.9.3) from Illumina. Sequencing data can be found within the NCBI BioProject PRJNA1007366.

### Bacteria and fungi community characterization

Paired-end raw sequencing reads were first trimmed by length and quality using Trimmomatic v.0.36, with options “LEADING:28 TRAILING:28 SLIDINGWINDOW:6:25 MINLEN:75” (48). Bowtie2 v.2.4.2 was used to map trimmed individual sample reads to bacteria and fungi marker gene databases, using the options “--no-discordant --very-sensitive” (49). The bacteria sequence database used was BEExact, a comprehensive 16S rRNA database of all bacteria taxa previously found associated with bees (50). The fungal sequence database used was SILVA 138.1 SSU, which contains 18S rRNA sequences (51). Samtools was used to transform mapping files and retrieve, with depth command, option “-a”, the coverage per base of the marker gene sequence (52). Coverage was summed for sequences from the same family or genus and normalized by the sequence length and library depth (*mean sequence coverage/normalization factor*; Table S1).

### Co-assembly and broad taxonomic assignment

Trimmed reads from all 18 samples were used in a co-assembly with MEGAHIT v.1.1.2 (53). Contigs >500 nt were binned into metagenome-assembled genomes (see below Metagenome-assembled bacteria genomes (MAGs)) and used for broad taxonomic classification of samples. We first classified bee contigs by mapping a 500 nt random fragment of contigs with Bowtie2 v.2.4.2 against *Apis mellifera* genome, GCF_003254395.2_Amel_HAv3.1 (49). Contigs that did not map were used as queries in a BLASTx (54) against nr database from NCBI, being classified according to the best hit taxonomy as bacteria, eukaryote/bee, virus or unknown (Table S2). Contig coverage used for the following analyses was recovered by mapping trimmed individual sample reads back to all contigs with Bowtie2 v.2.4.2, using the options “--no-discordant --very-sensitive” (49). Samtools was used for file transformation, and the depth command with option “-a” used to recover the coverage per base of the contigs (52).

### Metagenome-assembled genomes (MAGs)

Contigs were grouped into bins with MetaBAT2 v.2.11.3 using default settings (55). CheckM v.1.1.6 was used to assess the quality of the resulting MAGs (56). Bacterial MAGs with >30% completeness and <5% contamination were included in the following analyses. To confirm no need for dereplication of MAGs, which is expected for co-assemblies, we ran the “dereplicate” command from dRep v.3.4.2 (57). No MAGs were clustered beyond the threshold of >90% average nucleotide identity from the primary dendrogram of pair-wise Mash distances between all MAGs. Each MAG was subjected to gene/protein prediction with Prokka v.1.14.6, using default options (58). In addition, we used CRISPRCasFinder with default parameters to identify CRISPR-Cas loci in contigs from all MAGs (59). CRISPR repeat spacers from all CRISPR arrays were extracted and added to a spacer database used for putative phage host discovery (see “Characterization of bacteriophages in the microbiome”, List S1).

### Phylogenetic placement of MAGs and abundance

MAGs were placed into phylogenetic trees with related genomes for species-level taxonomic characterization. Best BLASTx results of the MAG contigs (see “Co-assembly and broad taxonomic assignment”) guided the choice of species reference genomes to retrieve from NCBI and include in the phylogenetic analysis, in addition to outgroups. Single-copy orthologs were recovered with OrhoFinder v.2.5.4 (60), aligned with MAFFT v.7.520 (61), and then concatenated for IQTREE v.2.2.0.3 maximum likelihood analysis, options -m TEST -B 1000 (62). Figtree v.1.4.4 was used for tree visualization. To estimate and compare MAG abundance across samples the coverage of single-copy orthologs was retrieved from previous mapping output (see “Co-assembly and broad taxonomic assignment”) with an in-house perl script (https://github.com/liliancaesarbio/general_scripts/), normalized to the sequence length and library depth (*mean sequence coverage/normalization factor*; Table S1).

### Microbiome functional characterization

All contigs classified taxonomically as bacteria, including MAG sequences, had proteins predicted with Prokka using default settings (63). The coverage of each protein was retrieved from previous mapping (see “Co-assembly and broad taxonomic assignment”) with an in-house perl script (https://github.com/liliancaesarbio/general_scripts/). Proteins were assigned for functionality at different levels of the KEGG database with EggNOG-mapper 2.1, using default parameters (64). Genes with >0.1 X coverage were considered present. First, KEGG level B of description was retried for each KO ID, and categories with total samples sum of >700 coverage and 100 protein counts were considered. Gene counts for both KEGG level B and KEGG level C for each non-general function category were plotted with R v.3.6.3, using package pheatmap (65). To test for functional similarity between samples the heatmap was hierarchically clustered with hclust complete linkage method.

### Characterization of bacteriophages in the microbiome

Sequences of bacteriophages were identified using three programs; geNomad was run with option “end-to-end” and a cutoff of >0.7 virus score (66), VirSorter2 v2.2.4 was run using options “--min-score 0.7 --hallmark-required-on-short” (67), and VIBRANT v1.2.1 was run using the default settings (68). For this analysis, contigs >500 nt, classified as bacteria, virus or unknown, were used as input, enabling the identification of lytic or lysogenic phages. For contigs of bacteria, only provirus (lysogenic phages) were considered, since the other detected phage sequences may represent horizontal gene transfer events. No viral bins were generated or dereplication needed due to the co-assembly approach (see Metagenome-assembled genomes (MAGs)”); also no bins were recovered with vRhyme v1.1.0, using default settings (69). CheckV “end-to-end” was used to estimate completeness of the final list of viral operational taxonomic units (vOTUs) (70). The coverage of vOTUs was recovered from the depth output files previously obtained (see “Co-assembly and broad taxonomic assignment”) using an in-house script (https://github.com/liliancaesarbio/general_scripts/). Coverage was estimated based only on the viral region of the contig since they are the result of a co-assembly, and in some samples that virus may not be present – as for the case of lysogenic phages. Candidate hosts were predicted with iPHoP v.1.3.1 (71), BLASTn with options “-evalue 1e-3 -ungapped -perc_identity 95” against NCBI nt database, CRISPRs spaces annotated from MAGs and also against the CrisperOpenDB (72). The last database includes more than 9 billion spacers, as well as the ones annotated in previously published bee microbiome studies (18, 44, 45).

### Queen genetic background

Trimmed reads from individual samples were mapped against *Apis mellifera* reference genome (GCF_003254395.2_Amel_HAv3.1) with Bowtie2 v.2.4.2, using the option “--no-discordant --very-sensitive” (49). Bam files were used to generate the beagle file of genotype likelihoods with ANGSD, options “-GL 2 -doGlf 2 -doMajorMinor 1 -SNP_pval 1e-6 -doMaf 1” (73). Since microbiome metagenome sequencing does not aim at a great sequencing depth of the host genome, here we follow the recommended approach of using genotype likelihoods to circumvent uncertainty of host genotypes due low or medium host sequencing coverage (74). Genetic structure and individual queen ancestry proportions were analyzed with NGSadmix, option -minMaf 0.05, for clusters (K) ranging from 2 to 5 (75). Population structure was further investigated by partitioning the genetic variance using a principal component analysis based on genotype likelihoods using PCAngsd (76). The contribution of each principal component (PC) was calculated in R and plotted using ggplot2 (77). As an alternative approach, used later for correlation analysis with microbiome composition, we also estimated pairwise genetic distances directly, using genotype likelihoods, with ngsDist (78). The input file for ngsDist was prepared with ANGSD, options “-minMapQ 20 -minQ 20 -doCounts 1 -GL 1 -doMajorMinor 1 -doMaf 1 -SNP_pval 1e-6 -doGeno 8 -doPost 1”.

### Statistical analysis

All graphs and analyses were run on R v.3.6.3. using the packages cited. We assessed beta diversity and clustering profiles for the 16S rRNA-based results with PCoA ordination plot ran on Bray-Curtis dissimilarity matrices using package Ape (79), then plotted with ggplot2 (77). To test for the effects of geographic location (*State* and *City*), year, age, queen origin, management, health status, and fitness markers (ovary size and ovary mass) on the bacterial community composition, we ran permanova, 9,999 permutations, with the function adonis2 from Vegan package (80). The effect of the same factors on the proportion of core microbiome members and Caudoviricetes was tested with *t-*test or Wilcoxon rank test and Kruskal-Wallis rank test in the case data was not normally distributed with Shapiro-Wilk normality test. All p-values were adjusted with the false discovery rate (FDR) method. To test for the co-occurrence of phages and bacteria we ran a Spearman correlation analysis with package Hmisc (81), plotted with packages corrplot (82) and PerformanceAnalytics (83). To test for the correlation between host genetic background and microbiome composition we used the Mantel test from package Vegan (80), with Spearman correlation and 9,999 permutations. The input matrices were the queen genotype likelihood covariance matrix or the queen pairwise genetic distance (see “Queen genetic background”), and the Bray-Curtis dissimilarity of the gut microbiome composition at the family or genus-level.

## Acknowledgements

LC was supported by the National Science Foundation IOS award #2005306, and by the GEMS Biology Integration Institute, funded by the National Science Foundation DBI Biology Integration Institutes Program, Award #2022049. AM’s work was supported by a L’Oreal-UNESCO For Women in Science Fellowship as well as a Project Apis m. grant, Boone-Hodgson-Wilkinson Trust grant, and a grant from the Eastern Apiculture Society. RU was supported by a USDA NIFA Award 2017-51300-26814.

---

**Figure S1:** Admixture analysis on genotype likelihoods inferring population structure of honey bee queens. Individual ancestry estimated for K3, K4 or K5 ancestral populations.

**Figure S2:** Maximum likelihood phylogenies of non-core bacteria present in the honey bee queen microbiome. Bifidobacteraceae inferred from the alignment of 289 single-copy orthologs (SCO), Enterobacteriaceae from 866 SCO, and Orbaceae from170 SCO.

**Figure S3:** Zoom-in on non-general function KEGG categories shown on figure 3 (main text). Color scale here is the absolute count of genes on that specific category.

**Figure S4:** Correlation plot of phage and metagenome-assembled genomes (MAGs) abundance. The color of circles corresponds to a positive (blue) or negative (red) Pearson’s correlation coefficient. The size of the circles corresponds to the p-value of the correlation coefficient. Only significant correlations (p < 0.05) are shown.

**List S1:** File with listed spacers from microbiome members.

**Table S1:** Metadata and sequencing statistics for each library from each queen sample.

**Table S2:** Contig statistics and taxonomic affiliation.

**Table S3:** Bin statistics, completeness, and contamination.

**Table S4:** Analysis of coverage for each bin across each queen microbiome sample.

**Table S5:** vOTUs recovered from the metagenomic samples and statistics on completeness and host prediction.

